# Dynamic optimism and pessimism states shape risk-taking behavior in monkeys

**DOI:** 10.64898/2026.05.01.722186

**Authors:** Iori Higashino, Ryo Ito, Yasushi Okochi, Kengo Inutsuka, Hiroshi Yokoyama, Rikako Kato, Yuichiro Yada, Ken-ichi Amemori, Honda Naoki

## Abstract

Humans and animals often face risky situations that require decision-making. Such decisions can be high-risk, high-return at some times, and low-risk, low-return at other times, depending on the balance between optimism and pessimism. However, how this optimism-pessimism bias is regulated across contexts remains unclear. Here, we introduced a computational model of decision-making in a risk-taking task based on the free-energy principle, together with a machine-learning framework that inversely estimates cognitive updating and optimism-pessimism bias from behavioral data. Applying this framework to monkey behavioral data, we found that a monkey quickly and accurately recognized the degree of risk, while frequently switching between optimism and pessimism during the task. In addition, we identified a characteristic control rule for optimism-pessimism bias that is distinct from reward-dependent regulation. Our framework provided a principled tool for understanding the latent cognitive processes underlying risky decision-making in animals and humans.

## Introduction

How do we make decisions when faced with risky situations? An important factor is the mental conflict between optimistic and pessimistic biases. For example, optimistic individuals tend to pursue high-risk, high-return options^1^, whereas pessimistic individuals prefer low-risk, low-return alternatives^2^. Both extremes can be considered unreasonable for maximizing expected reward. Importantly, this bias is not static; for instance, optimism and pessimism vary within individuals over short timescales.^3^ Moreover, the temporal dynamics of optimism-pessimism (OP) bias are closely linked to psychiatric symptomatology, especially depression. In young adults, intra-individual fluctuations in state optimism over weeks are associated with changes in depressive mood^4^. These observations underscore the importance of OP bias in risky decision-making in uncertain environments, yet no established method exists to quantify its temporal dynamics.

Decision-making has recently been modeled within the framework of the Free Energy Principle (FEP) under the Bayesian brain hypothesis^5–11^. Moreover, both FEP and RL are grounded in the assumption of rational optimization, minimization of free energy (or maximization of information gain) in FEP, and maximization of expected reward in RL. While extensions of FEP can represent optimism and pessimism through preferred outcomes or asymmetric weighting of prediction errors, these factors are generally treated as stationary within a task. Reinforcement learning (RL) has similarly been expanded to capture optimistic and pessimistic choice tendencies^12,13^, but the degree of OP bias in these models is likewise fixed rather than dynamic. Consequently, existing theoretical frameworks remain limited in explaining within-individual, time-varying OP bias, and there is currently no established method to quantify its temporal dynamics. By contrast, our aim was to capture non-rational decision-making, focusing on the temporal dynamics of optimism-pessimism (OP) bias that deviate from purely optimal strategies.

An important next step is to quantify the temporal dynamics of OP bias in order to investigate its neural basis. Many computational studies have been devoted to constructing theories of decision-making, but they have largely neglected the temporal dynamics of internal states^14,15^. Thus, there is a strong need for methods that can decode such time-varying internal states directly from behavioral data (**Fig. 1a**). Developing such approaches would make it possible to analyze the neural correlates of the temporal variability of optimism and pessimism, thereby providing insights into how the brain regulates OP bias in a context-dependent manner.

**Fig. 1:**
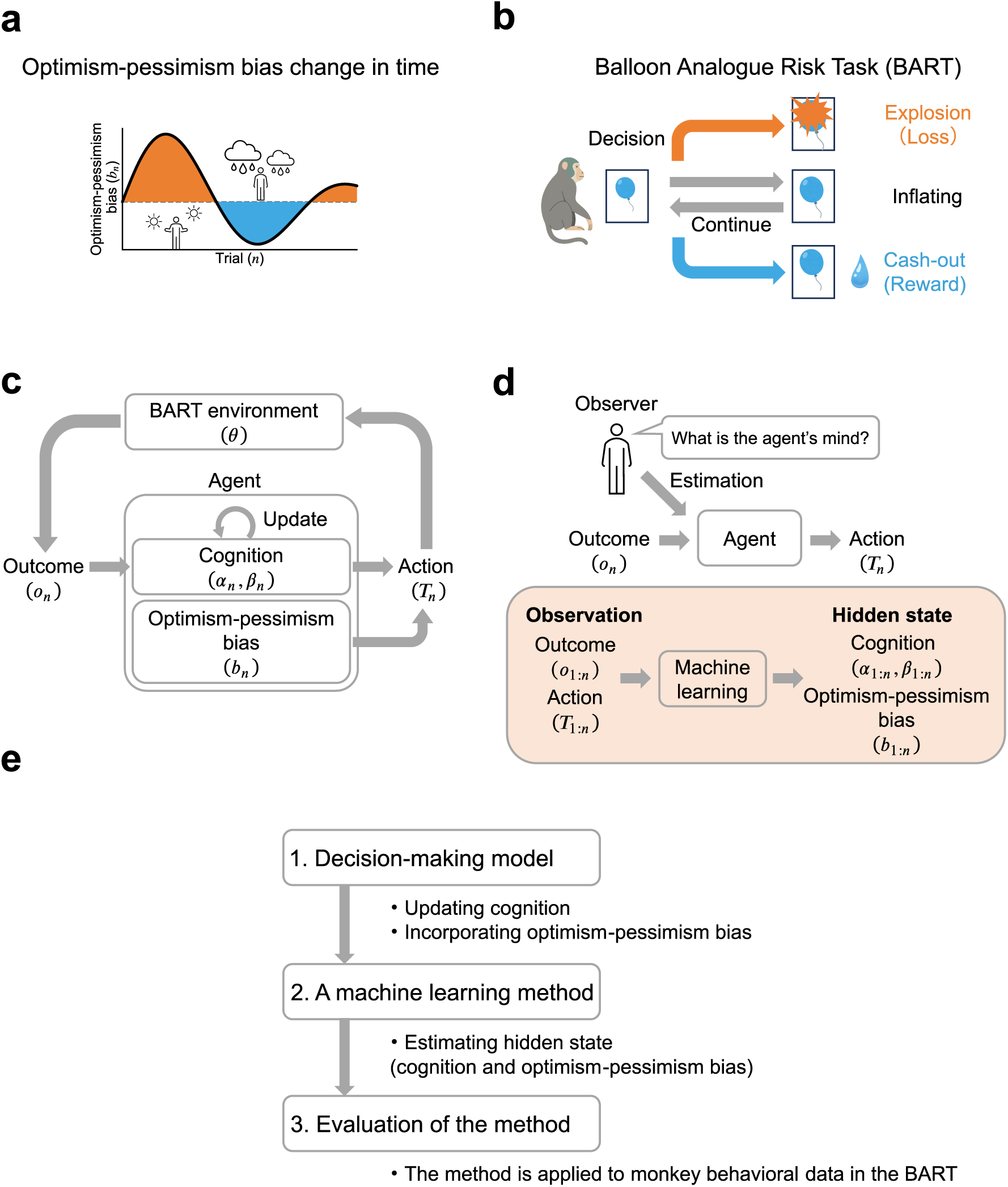
Overview of the framework for decoding dynamic optimism-pessimism bias. (**a**) The temporal dynamics of optimism-pessimism bias (*b_n_*) in risk-taking behavior. (**b**) Scheme of the BART. The balloon linearly inflates over time and explodes at a random time point. If the monkey cashes out before the explosion, the monkey can receive a reward. (**c**) The computational model of the decision-making agent (Dynamical OP model). The agent updates its internal cognition (*α_n_* and *β_n_* are the shape and rate parameters of gamma distribution, respectively) about the BART environment (*θ*) from the outcome (*o_n_*) and selects action (*T_n_*) with OP bias (*b_n_*). (**d**) A machine learning method (called inverse OP method) to estimate dynamical changes in hidden states from observed time-series data, from the viewpoint of an observer of the agent. (**e**) Flow of this study. 1. Formulating the agent’s decision-making model (dynamical OP model). 2. Developing a machine learning method to estimate hidden states of the agent based on the dynamical OP model. 3. The method is applied to monkey behavioral data in the BART.

To address this issue, we focused on the Balloon Analogue Risk Task (BART)^16^, a well-established experimental paradigm for assessing risk-taking behavior in humans (**Fig. 1b**). The BART was particularly useful for examining behavior associated with dynamic fluctuations in optimism-pessimism (OP) bias. We then developed a computational model of decision-making in the BART based on FEP, with OP bias represented as a latent state that modulates risk-taking behavior (**Fig. 1c**). Based on this model, we further developed a machine-learning method to infer the agent’s OP bias and risk cognition from monkey behavioral data in the BART (**Fig. 1d**). We thereby identified a characteristic rule governing the dynamics of OP bias that differs from simple reward-dependent regulation.

## Results

### Decoding optimism and pessimism in risk-taking behavior

We first presented the overall framework of this study (**Fig. 1**). The goal was to infer time-varying OP bias (**Fig. 1a**) from behavioral data in the BART. In the BART, an agent observed a continuously inflating balloon whose size corresponded to accumulated reward and had to decide whether to keep waiting for a larger reward or cash out before the balloon exploded. In these repeated trials, decision-making could be dynamically affected by a conflict between optimistic risk seeking and pessimistic risk avoidance (**Fig. 1b**).

Our framework consisted of three steps. First, we formulated a decision-making model in which the agent updated risk cognition and selected actions under OP bias (**Fig. 1c**). Second, we developed a machine learning method called the inverse OP method that estimated latent OP bias and risk cognition from observed actions and outcomes (**Fig. 1d**). Third, we applied this method to monkey behavioral data in the BART for characterizing how OP bias evolved over trials and across days (**Fig. 1e**).

In the following section, we will describe the details for each of the three-step procedures and describe how these procedures are linked and integrated to establish our proposed framework.

### Dynamical OP model

We formulated a computational model of the dynamic decision-making process in the BART, where optimism and pessimism biases fluctuated across trials. This model, called the Dynamical OP model (DOP model), consisted of two distinct processes involved in decision-making under risky situations. Firstly, we considered a cognitive process, in which the agent updated its belief about the rate of increase in explosion probability as the balloon inflated, based on Bayesian inference (**Fig. 2a**). Secondly, we defined an action process, in which the agent decided whether to continue inflating or to cash out the accumulated reward, depending on its current belief and OP bias (**Fig. 2b**). The agent repeated these two processes throughout the BART.

**Fig. 2:**
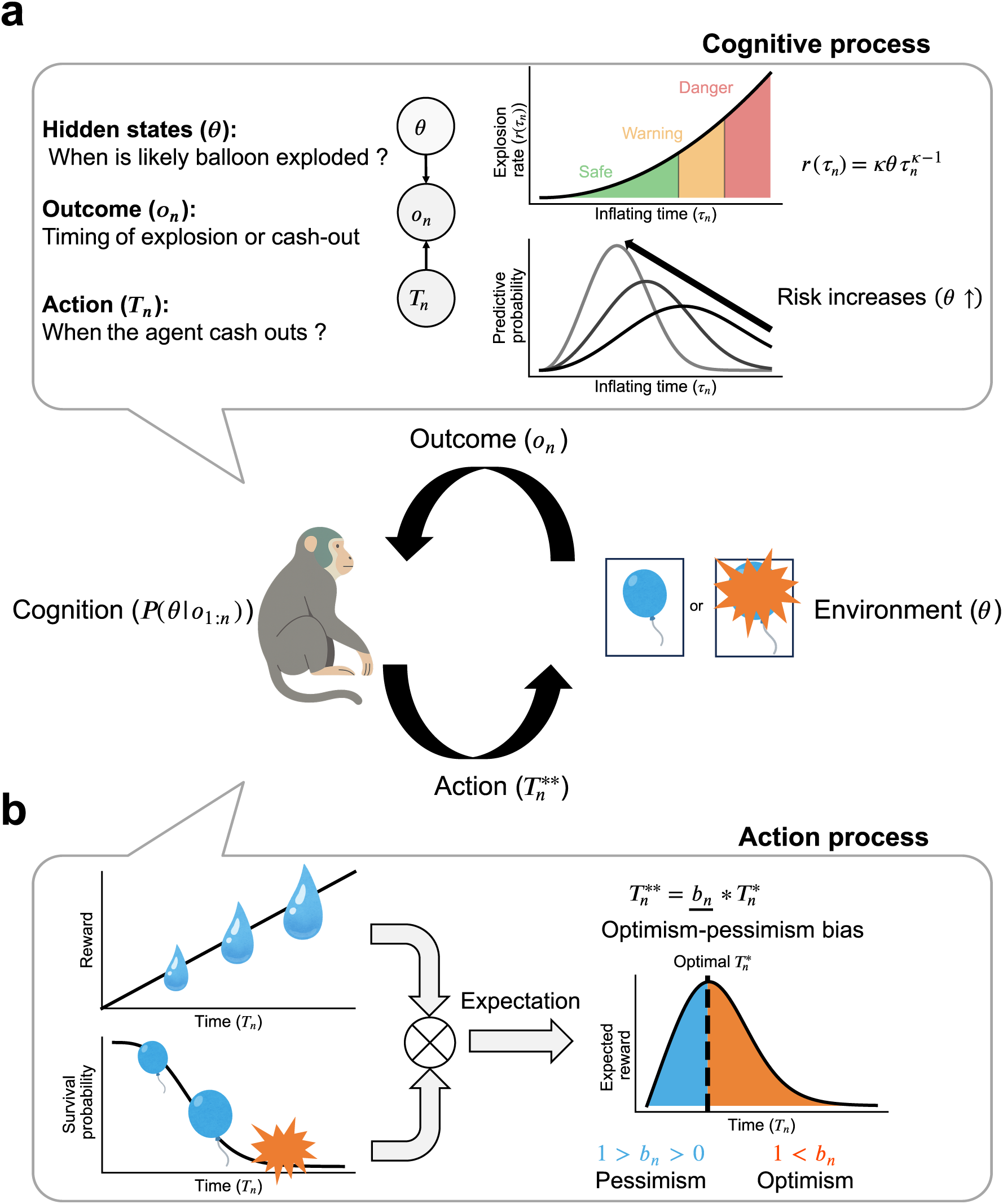
Decision-making model for the Balloon Analogue Risk Task (BART) with optimism-pessimism bias. (**a, b**) The computational framework of decision-making in the BART. The agent (the monkey) interacts with the environment in the BART by repeating cognitive and action processes. *P*(*θ*|*o*_1:*n*_) is a cognitive distribution of environmental risk *θ* given observed timing of explosion or cash-out *o*_1:*n*_ from 1 to *n*. (**a**) Cognitive process. The monkey assumes that given cash-out timing, either explosion or not is probabilistically observed from the explosion rate (**Left**). The explosion rate increases over time as 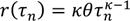, where *θ* is a hidden parameter controlling how fast the explosion probability inflates (**Upper right**). Through trial and error, the monkey infers *θ* for each trial and updates the cognition of when the balloon likely explodes (as predictive probability of explosion timing). (**b**) Action process under optimism-pessimism bias. The monkey evaluates the expected reward based on the product of the amount of reward (Upper left panel) and survival probability of the balloon (**Lower left**). The optimal action is to cash out the reward at the timing that maximizes the expected reward. The action timing is modulated by the optimism-pessimism bias (**Right**).

In the cognitive process, the agent internally had a world model in which either explosion or not was probabilistically observed from the explosion rate (**Fig. 2a, left**), and the explosion rate nonlinearly increased over time (**Fig. 2a, upper right**). Based on this world model, the agent sequentially estimated the parameter *θ* using Bayesian updating; as *θ* increased, the expected explosion timing became earlier, resulting in a horizontal shift of the predicted explosion-time distribution (**Fig. 2a, lower right**). Note that the predicted explosion-time distribution was parametrized by *α_n_* and *β_n_*, which controlled the explosion-risk growth rate and the update of these parameters was formulated by the Free Energy Principle (FEP) (**see Methods in detail**).

In the action process, the agent decided whether to continue inflating or to cash out based on its current belief about the explosion risk, as updated in the cognitive process. The agent was guided by two opposing factors. The first factor was that waiting longer increased the accumulated reward (**Fig. 2b, upper left**). The second factor was that waiting longer also increased the risk of explosion (**Fig. 2b, lower left**). The agent determined the optimal timing of cash-out 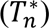 by balancing these two opposing factors.

We introduced the meta-parameter of OP bias, *b_n_*, which modulated the timing of decision-making, where *n* indicated the trial (**Fig. 2b right**). When the bias became optimistic, the agent tended to wait longer than the optimal timing, thereby taking greater risks. Conversely, when the bias became pessimistic, the agent cashed out earlier and avoided risk. When the bias was neutral, the agent behaved optimally. Note that we assumed dynamic mental conflict between optimism and pessimism as *b_n_* varied over trials.

### Simulation of DOP model

To validate the model, we simulated an agent performing the BART, where explosion timings were randomly sampled across trials (**Fig. 3a, b**). For neutral OP bias (i.e., *b_n_* = 1), the agent alternated between cashing out and experiencing explosions (**Fig. 3b**). From these observed outcomes (i.e., whether the balloon exploded or the agent successfully withdrew), the model agent inferred the predictive distribution of explosion timings, which gradually converged toward the ground truth distribution (**Fig. 3a, upper**). However, the predicted distribution was slightly shifted toward earlier times, indicating that the agent expected the balloon to explode earlier than the average true explosion time. As a result, the agent tended to cash out earlier than optimal, producing a small deviation from the ground-truth distribution. From the predicted explosion-time distribution, we calculated the survival probability of the balloon *S*(*τ_n_*), which represented the probability that the balloon had not exploded by time *τ_n_*. *S*(*τ_n_*) remained high at early times and dropped sharply around ∼2 s (**Fig. 3a, middle**). We also computed the hazard function ℎ(*τ_n_*), defined as the instantaneous rate of decline in survival probability normalized by its current value 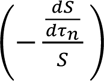. The hazard function was initially low, increased between 2 and 6 s, and then decreased thereafter (**Fig. 3a, bottom**).

**Fig. 3:**
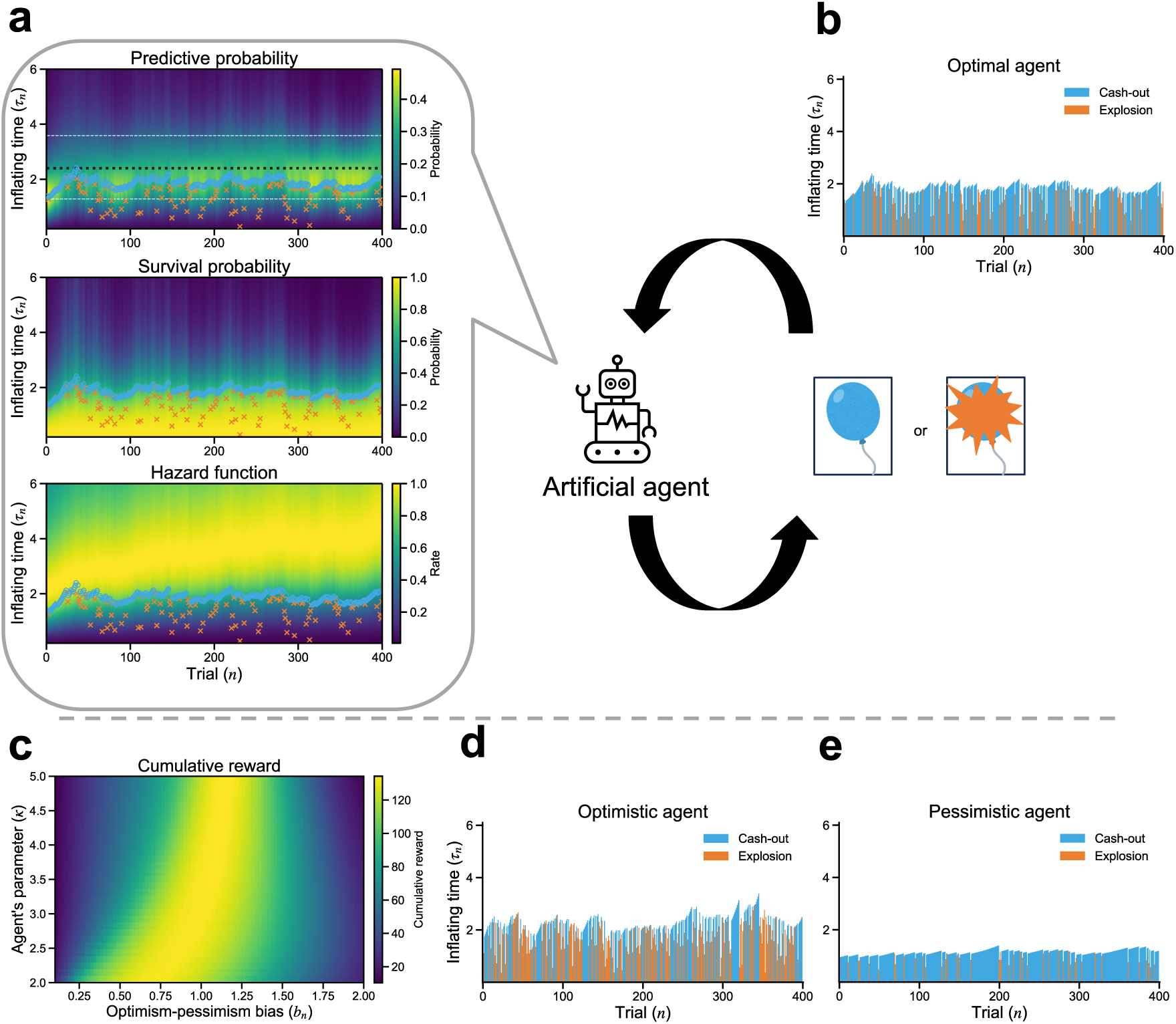
Simulation of decision-making model in the BART. (**a, b**) Simulation of an artificial optimal agent (*b_n_* = 1). (**a, upper**) Predictive probability of explosion timing as a time-by-trial heat map. Red cross and blue circle indicate timings of explosion and cash-out, respectively. Black and white dashed lines indicate median and 10%/90% quantiles of true explosion probability. (**a, middle**) A heat map of survival probability of the balloon estimated by the agent. (**a, lower**) A heat map of hazard function. (**b**) Cash-out/explosion timings of the agent. (**c**) Cumulative reward depending on optimism-pessimism bias. Cumulative reward is computed excluding the learning phase (the 100 trials). (**d, e**) Cash-out/explosion timings of the optimistic and pessimistic agents, respectively (*b_n_* = 1.3 (**d**) and *b_n_* = 0.7 (**e**)).

In the simulations so far, we assumed that the agent knew the ground truth of the explosion risk growth (i.e., *κ*). We then relaxed this assumption. We performed simulations where the explosion risk grew with *κ*^∗^ = 3, while the agent held a misspecified belief about the shape parameter *κ*. Varying the believed *κ* as well as the OP bias, we computed cumulative reward (**Fig. 3c**). When the agent overestimated the explosion risk growth (i.e., *κ* > 3), the optimal bias was consistently close to 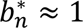. Then, the agent’s predicted distribution of explosion times closely matched the true distribution (**Supplementary Fig. 1**). In contrast, when the agent underestimated the explosion risk growth (i.e., *κ* < 3), the optimal *b_n_* became biased toward smaller. These results indicated that the DOP agent that overestimated the explosion risk growth correctly recognized the explosion timing structure, which indicated that neutral OP (i.e., *b_n_* ≈ 1) was optimal in terms of reward maximization.

### Optimistic and pessimistic behaviors in the model

Next, we examined how the OP bias modulated risk-taking behaviors driven by optimism or pessimism. First, we performed simulations with optimistic bias (*b_n_* = 1. 3) and observed that the agent preferred high-risk, high-return actions. This optimistic agent frequently caused balloon explosions and thus rarely obtained large rewards (**Fig. 3d**). The agent continuously selected long cash-out times, resembling the behavior of adolescent cannabis users who pursue high rewards despite risky environments^17^. Second, we performed simulations with pessimistic bias (*b_n_* = 0.7) which led to the agent preferring low-risk, low-return actions. This pessimistic agent avoided explosions entirely, repeatedly made early cash-outs, and consequently accumulated small but reliable rewards (**Fig. 3e**). Such seemingly irrational behavior reflected a preference for certain rewards (i.e., low-risk, low-return) and parallels the risk-avoidant tendencies observed in patients with obsessive-compulsive disorder (OCD)^18^. Together, these simulations demonstrated that varying the OP bias enabled our model to capture both optimistic and pessimistic action processes, producing corresponding high-risk, high-return and low-risk, low-return behaviors.

### Inverse Optimism-Pessimism method (inverse OP): Estimation of the OP bias

In the above cases, we assumed a constant OP bias. However, our internal states, e.g., optimism and pessimism, varied in a context-dependent manner. Quantitative evaluation of temporal variations in the agents’ hidden state was important for elucidating their neural mechanisms. Here, we used a machine learning approach called inverse OP^10^ to estimate dynamical OP bias parameters. In inverse OP, we adopted a shift in perspective from the agent to the observer of the agent. We first addressed the sequential cognition of explosion probabilities by the agent. Then, we addressed the estimation of the hidden state of the agent by the observer of the agent (**Fig. 1d, 4a**). To this end, we developed a state-space model (SSM) which described the dynamics of latent variables (OP bias and cognition) and the generative process of observations (i.e., cash-out or explosion timing). Leveraging this SSM, we sequentially estimated the hidden state of the agent (*z*) from observations (*o*) in a Bayesian manner:

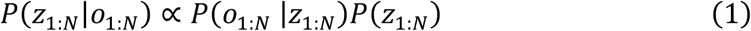

where *o_n_* = {*δ_n_*, *τ_n_*}, *z_n_* = {*b_n_*, *α_n_*, *β_n_*} and the subscript 1: *N* indicated steps 1 to *N*. This Bayesian estimation, namely inverse OP, was conducted using a particle filter and particle smoother (**See Methods for details**).

To test the validity of the inverse OP method, we applied it to the artificial data generated by the DOP model. We simulated a model agent with nonconstant OP bias in the BART. We then demonstrated that inverse OP estimated the ground truth hidden state of the simulated agent, that is, the agent’s intensity of OP bias and cognition (**Supplementary Fig. 2**). Thus, inverse OP provided efficient estimators of temporal fluctuations in the conflict between optimism and pessimism.

### inverse OP decoding of OP bias behind monkey behavior

We applied inverse OP to monkey behavioral data in the BART, in which explosion timings were uniformly sampled from 2 s to 6 s. The monkey tended to cash out early in the initial phase but later delayed cash-out, resulting in more explosions while the overall reward acquisition rate increased (**Fig. 4b**). Using inverse OP, we found that the monkey’s OP bias changed dynamically over the trials: it was pessimistic in the early phase (0 < *b_n_* < 1) and gradually shifted toward a neutral state (*b_n_* ≈ 1) in the later phase (**Fig. 4c**). This smooth transition in the decoded OP bias was consistent with the observed behavioral shift from risk avoidance to increased risk taking. We also successfully estimated how the monkey recognized and updated the predictive distribution of explosion timing on a trial-by-trial basis (**Fig. 4d**). The peak of the predicted distribution approached the mean of the uniform distribution, around 4 s, indicating that the monkey correctly learned and recognized the environmental statistics. In addition, we visualized the survival probability *S*(*τ_n_*) (**Fig. 4e**) and the hazard function ℎ(*τ_n_*) (**Fig. 4f**), indicating that the monkey took cash-out action around the time when the survival probability suddenly dropped and hazard function started to increase. Taken together, these results demonstrated that inverse OP quantitatively captures the monkey’s OP bias and risk cognition dynamics.

**Fig. 4:**
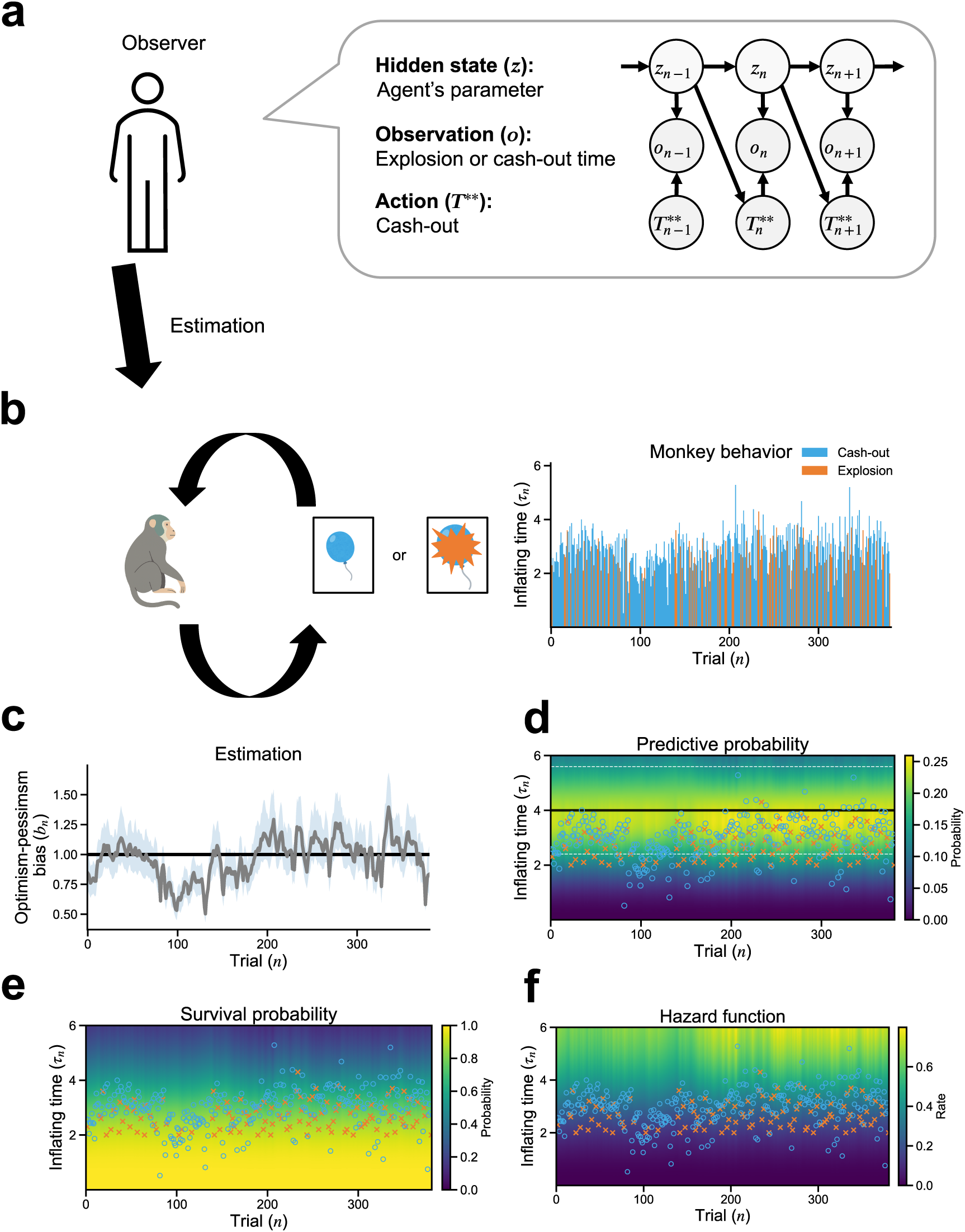
inverse OP decoding temporal dynamics of OP bias in a monkey performing BART. (**a**) Estimation of the hidden internal state of the monkey by observing the monkey’s behavior in the BART. The graphical model (right) represents state-space model from the perspective of an observer of the monkey. The OP bias (*b_n_*), and parameters representing the monkey’s cognition (*α_n_*, *β_n_*) are the hidden variables, whereas the outcome (*o_n_*) and action 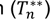 are observed variables for the observer. (**b**) Cash-out/explosion timings of the monkey. (**c**) inverse OP-decoded OP bias over trials: posterior mean (solid line) with 95% credible band (shading, 2.5%-97.5%). The horizontal solid line marks the unbiased/optimal reference (i.e., *b_n_* = 1). (**d-f**) Estimation of the monkey’s internal cognition. The predictive probability of explosion timing (**d**), survival probability of the balloon (**e**), and hazard function (**f**).

### Reward acquisition-driven dynamics of OP bias across days

We next examined decision making across days. The monkey performed the BART over eight days (with the inter-day intervals shown in the figure). The decoded OP parameter exhibited day-to-day modulation: OP bias was high on day 1 (*b_n_* > 1), dropped on day 2 (*b_n_* < 1), and subsequently converged toward values around 1 over the remaining days (**Fig. 5a**). This temporal pattern was well aligned with the observed behavioral changes. On day 1, reward acquisition was high, consistent with high-risk, high-return behavior driven by an optimistic OP bias (*b_n_* > 1). Correspondingly, the proportion of explosion trials was also high. On day 2, reward acquisition markedly decreased, reflecting a shift toward low-risk, low-return behavior associated with a pessimistic OP bias (*b_n_* < 1), accompanied by a reduced proportion of explosion trials (**Fig. 5b**). Across days 3-8, reward acquisition gradually increased as OP bias stabilized around the neutral point (*b_n_* ≈ 1) (**Fig. 5c**). During this period, the proportion of explosion trials also showed a gradual increase but declined again during the final two days. This pattern suggested that the monkey first engaged in optimistic exploration, then shifted to a more conservative, pessimistic mode, and later approached a more balanced, less biased policy. These findings showed that OP bias varied across days and was associated with changes in risk-taking behavior. We next examined what factors drove this temporal modulation of OP bias.

**Fig. 5:**
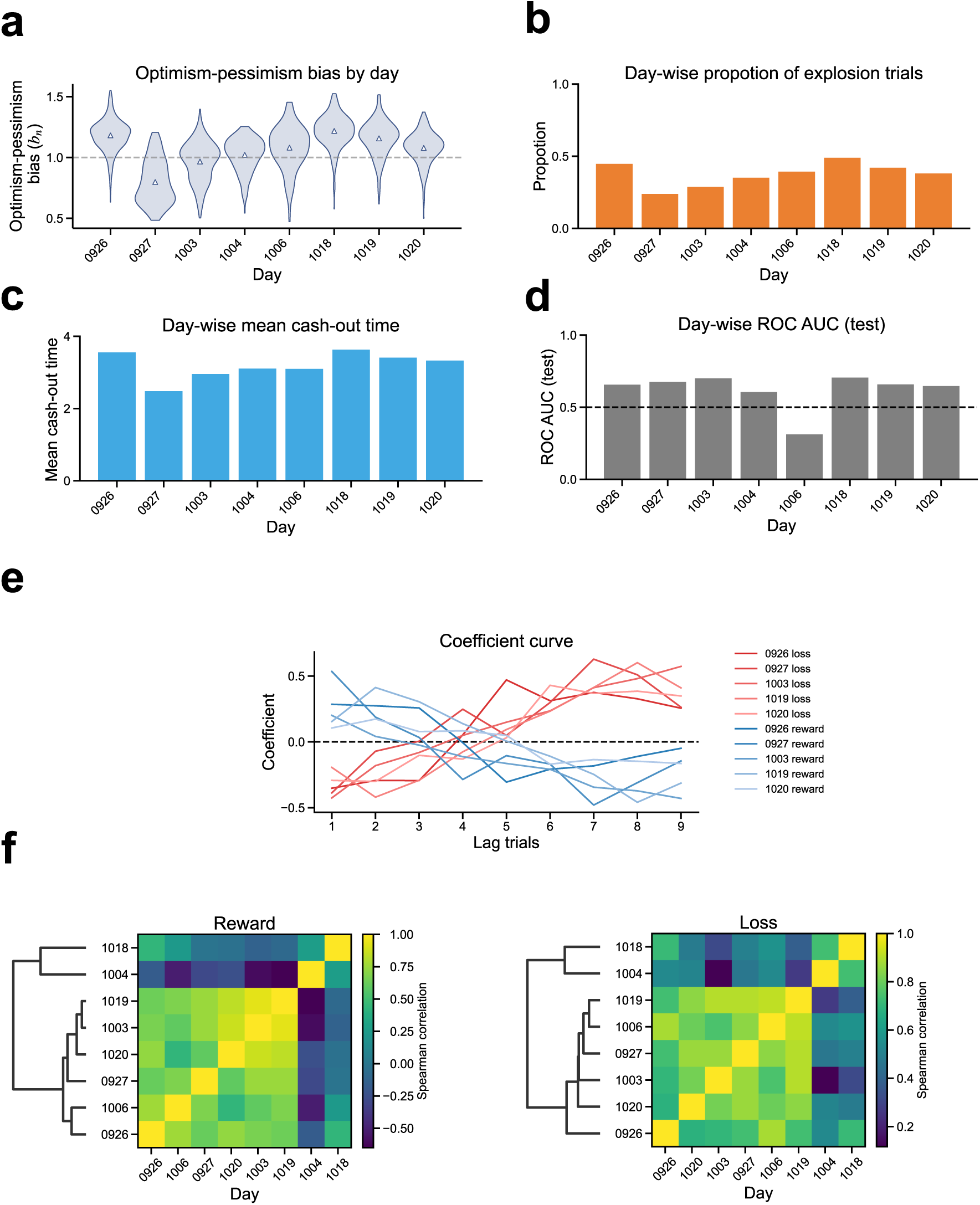
OP bias is driven by explosion and cash-out time. (**a**) Day-wise distribution of the decoded OP-bias parameter *b_n_* (violin plots; triangles indicate the mean). The dashed line marks the unbiased reference level (*b_n_* = 1). Numbers on the x-axis denote the date of the experiment (0926 means September 25). (**b**) Proportion of explosion trials for each day. (**c**) Mean cash-out time for each day. (**d**) Performance of the logistic classifier predicting either up or down of the OP bias estimation from the amount of reward and loss for the past nine trials. Performance is evaluated by ROC AUC. Dashed line indicates chance level, 0.5. (**e**) Lag-wise coefficients of the logistic classifier. Blue and red lines indicate coefficients for reward and loss, respectively. (**f**) Cross-day similarity of the lag-wise logistic classifier coefficients for reward and loss, quantified by Spearman correlation (color scale). Dendrograms are based on hierarchical clustering with correlation distance (1 − *Spearman correlation*).

To address this question, we examined how OP bias was regulated by recent experiences of reward acquisition and loss. We formulated a classification problem to predict whether OP bias would increase or decrease on the next trial based on the history of reward acquisition and loss over the preceding nine trials. This problem was solved each day using logistic regression:

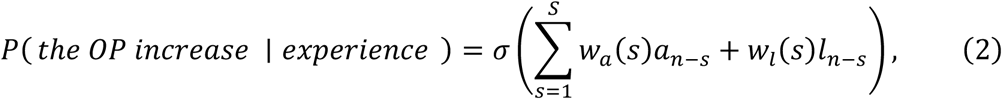

where *σ*(*x*)=1/(1+exp(-*x*)); *a_n_*_–*s*_ and *l_n_*_–*s*_ indicate reward and loss at lag *s*, respectively; *w_a_*(*s*) and *w_l_*(*s*) indicate the lag-dependent coefficients that quantified the contributions of reward acquisition and loss to OP increases. The coefficients *w_a_*(*s*) and *w_l_*(*s*) were estimated for the first half trials of each day by the maximum likelihood method.

On most days (with day 5 as an exception), the model classified OP increases versus decreases modestly above chance (AUC of test data (last half trials of each day) ≈ 0.6-0.7) (**Fig. 5d**), suggesting that OP dynamics were influenced by recent reward and loss experiences. To characterize day-specific OP control patterns, we clustered the estimated coefficients across days (**Fig. 5f**). This analysis revealed one major cluster comprising days 1-3, 5, and 7-8 (major coefficients cluster), as well as two outlier profiles observed on days 4 and 6 (minor coefficients cluster, **Supplementary Fig. 3**).

Within the major coefficients cluster, although we initially expected reward losses to dominantly drive OP regulation, the estimated coefficients *w_a_*(*s*) and *w_l_*(*s*) instead exhibited nearly symmetric profiles (**Fig. 5e**). This symmetry indicated that reward acquisition and loss exerted comparable influences on OP dynamics, such that OP was regulated by the overall amount of recent outcomes rather than by a qualitative distinction between reward acquisition and loss. Moreover, the RF profiles showed that very recent outcomes tended to promote increases in OP (greater optimism), whereas outcomes further in the past tended to promote decreases in OP (greater pessimism). These results suggested that OP control was not well captured by a purely reward-driven learning account.

In contrast, the minor coefficients did not exhibit symmetric profiles. For these days, early lags contributed little to OP modulation, whereas more distant lags were associated with decreases in OP. This distinct temporal structure suggested that qualitatively different OP control mechanisms may have been engaged on these days.

## Discussion

Decision making under risk has long been studied, particularly the choice between a high-risk, high-return (optimistic) strategy and a low-risk, low-return (pessimistic) strategy. However, most previous studies have treated the balance between optimism and pessimism as a time-invariant, stable trait^11–13^. In contrast, we conceptualized this balance as a context-dependent, temporally evolving state that can dynamically change over time. To this end, we presented inverse OP, a data-driven approach to decode time-varying optimism-pessimism (OP) bias directly from behavioral data. This approach provided a quantitative way to link temporally fluctuating biases to underlying neural mechanisms, as well as to clinically relevant instabilities in decision making.

Applying inverse OP to monkeys’ behavioral data in the BART, we successfully decoded temporal dynamics of OP bias. Notably, optimism and pessimism alternated even within a single day, suggesting that OP bias reflects experience-dependent state changes rather than a fixed trait. Moreover, although OP trajectories varied across days, repeated task experience appeared to drive OP toward an adaptive value near optimality (*b* ≈ 1). This pattern implied the possibility that monkeys gradually learned to regulate OP itself through experience, potentially reflecting a form of meta-learning over internal bias. Finally, we found a characteristic control rule for OP dynamics: OP was modulated by recent outcomes in a manner that depended on the overall magnitude of reward acquisition and loss, rather than on their valence. In other words, positive and negative outcomes exerted comparable influences on OP regulation. This result suggested that OP control may not be fully explained by a purely reward-driven learning account and instead may involve a more general meta-learning-like process.

Dynamic OP bias might resemble mood-related fluctuations observed in psychiatric disorders. For example, major depressive disorder may be characterized by persistently pessimistic states, whereas bipolar disorder involves alternating manic and depressive episodes that could correspond to switches between optimistic and pessimistic modes of decision making^19^. Similarly, individuals with gambling disorder or substance use disorder often exhibit distorted risk evaluation, which could be interpreted as forms of excessive optimism^17,20^. Across these conditions, a common feature was a dysregulated balance between optimism and pessimism, accompanied by impaired control over risky decision making.

To date, much effort in psychiatric research has focused on identifying biomarkers that characterize static traits associated with mental disorders^21–26^. In contrast, biomarkers that track within-subject temporal variations provide a complementary perspective by quantifying fluctuations in optimism-pessimism bias over time. Such dynamic markers may also offer quantitative targets for intervention, for example by promoting states closer to an optimal balance (e.g., *b_n_* = 1). In this way, our approach would contribute to the identification of biomarkers that capture moment-to-moment variability in mental states, rather than fixed pathological traits.

Several mathematical models have been proposed to explain decision making in the BART. A seminal theoretical account was provided by Wallsten et al.,^27^ who developed a model combining Bayesian inference of explosion probability with prospect-theoretic decision rules. By fitting the model to human BART data, they showed that participants’ behavior was best explained by assuming a stationary belief about explosion probability and a pre-determined action policy (i.e., when to stop pumping). This framework has since been extended in several variants^28,29^, further improving the descriptive accuracy of BART behavior. Conceptually, these models assume Bayesian-optimal belief updating in discrete time and do not explicitly incorporate OP bias. In contrast, our approach departed from Bayesian optimality by adopting a FEP-based formulation, operated in continuous time, and explicitly incorporated optimism-pessimism bias. More recently, pessimism-related parameters have been introduced into extensions of the Wallsten framework^30,31^. In these models, pessimism is implemented at the level of belief updating, such that prior beliefs are weighted more strongly than observational likelihoods, leading to pessimistic estimates of explosion probability. While these studies focus exclusively on pessimism and assume time-invariant parameters, our model introduced both optimism and pessimism and allowed their balance to vary dynamically over time. Other extensions of the Wallsten model have proposed alternative utility functions that incorporate reward variance in addition to expected reward and loss. Within such formulations, avoidance of reward loss can be interpreted as a form of pessimism, whereas preference for high reward variance may be viewed as optimism. However, these interpretations remain implicit, and the associated parameters are typically assumed to be fixed over time. In contrast, our approach explicitly modeled optimism-pessimism bias as a temporally evolving latent state and estimated its dynamics directly from behavioral data. Taken together, whereas existing models of BART behavior assumed Bayesian-optimal agents and, when optimism-pessimism-related factors were included, treated them as time-invariant, our work modeled non-optimal Bayesian agents to capture gradual learning and represent a time-varying OP bias that is inferred in a fully data-driven manner.

Our formulation builds on the free-energy principle. Whereas our recent inverse-FEP work focused on decoding time-varying trade-offs between reward seeking and curiosity in bandit tasks, the present study shifts attention to a risk-related mental conflict: the dynamic balance between optimism (positive expectations about future outcomes) and pessimism (negative expectations about future outcomes). We introduced a data-driven method to infer this latent optimism-pessimism (OP) dynamics directly from behavior. Conceptually, this framework generalized inverse approaches from inferring conflicts over what to pursue to inferring conflicts over how outcomes are evaluated under risk and uncertainty. Importantly, this line of work is grounded in the view of bounded rationality, in which human and animal decision makers operate under limited computational and informational resources^32,33^. In parallel, a broad class of inverse modeling approaches has been developed to estimate internal states from behavioral data, including Bayesian models of the BART^27^, reinforcement learning, and FEP or so-called meta-Bayesian approaches that explain individual differences and contextual dependence through inference over latent states^34–37^. However, these models typically assume static internal states or fixed parameters, which limits their ability to capture the evolving nature of real-world decision making in humans and animals. In contrast, our approach provided a computational account of decision making that explicitly incorporates bounded rationality in the form of time-varying internal states.

Our framework represented a first step toward quantifying time-varying optimism-pessimism (OP) from BART behavioral data, and several simplifying assumptions point to clear directions for future refinement. We assumed that decisions were fully formed before each trial; however, agents may change their minds as the balloon inflates. Capturing such within-trial hesitation will require formulations that incorporate two coupled state-space models, operating both across trials and within individual trials. Moreover, modeling the dynamics of the latent OP state itself offers a natural direction for extension. In the current formulation, OP evolves as a random walk; however, our results indicated that OP fluctuations are systematically driven by experience (**Fig. 5e**), suggesting context-dependent regulation. Introducing drift terms reflecting both internally dependent dynamics and externally input-dependent control into the OP evolution, and estimating these control functions directly from data, would enable explicit characterization of the mechanisms governing shifts between optimistic and pessimistic modes. Finally, linking the inferred OP dynamics to neural activity would help identify the neural substrates underlying OP bias and open avenues for targeted intervention. Together, these directions outlined a unified framework for elucidating the context-dependent control principles of OP and their neural implementation.

## Methods

### Generative model for cognition of explosion timing

In the BART, the agent assumes that balloon bursts are probabilistically generated by an inhomogeneous Poisson process with a time-increasing rate:

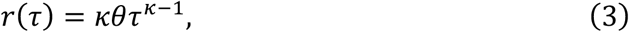

where *τ* denotes the explosion time, *κ* is a nonlinearity (shape) parameter, and *θ* controls the overall scale of risk. Under this assumption, the probability density function of the waiting time until the first explosion follows a Weibull distribution^38^:

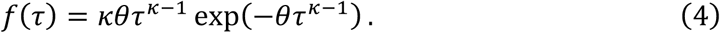

Note that the Weibull distribution is widely used in survival analysis to model time-to-event data, such as failure times of mechanical or electronic devices and component lifetimes.

### Agent’s evaluation of outcomes

In the BART, the agent observes one of two event types on each trial: explosion or cash-out. Each outcome is characterized by the event timing *τ*. Under the generative model above, the probability of explosion timing follows a Weibull distribution,

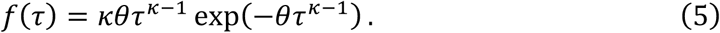

The probability of cash-out timing τ is given by

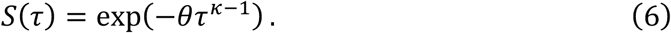

Note that *S*(*τ*) is known the survival probability, which can be derived by 1 – 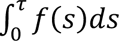. Because each BART trial results in either an explosion or a cash-out, the likelihood of an outcome can be written compactly as

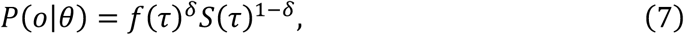

where *δ* ∈ {0,1} indicates the event type (*δ* = 1 for explosion, *δ* = 0 for cash-out), and *o* = {*τ*, *δ*} denotes the observed outcome.

### Optimal Bayesian update model for explosion rate

Based on the generative model of explosion timing (Eq. (7)), we assume that the agent recognizes the increasing rate of explosion by inferring the parameter *θ*. This cognitive process is modeled as Bayesian belief updating across trials. The posterior distribution over *θ* is updated according to Bayes’ rule:

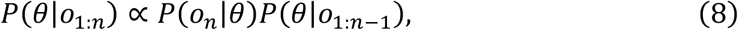

where *n* denotes the BART trial index and *o*_1:*n*_ denotes the sequence of outcomes up to trial *n*. *P*(*o_n_* ∣ *θ*) is the likelihood of the current outcome, and *P*(*θ* ∣ *o*_1:*n*–1_) is the prior belief carried over from the previous trial.

The prior distribution over *θ* is assumed to follow a Gamma distribution,

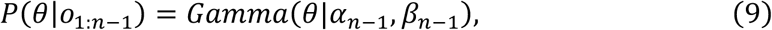

where *α_n_* and *β_n_* denote the shape and rate parameters, respectively. Importantly, in this Bayesian update scheme, the Gamma distribution serves as a conjugate prior for the Weibull likelihood (or equivalently, for the associated survival probability in the case of cash-out), so that the posterior distribution over *θ* remains a Gamma distribution.

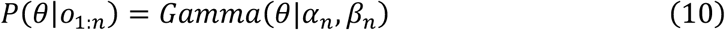

Then, the posterior parameters are updated as

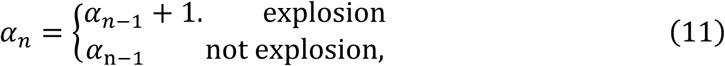

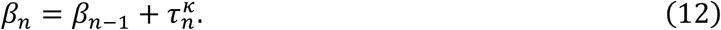

### Sub-optimal Bayesian update based on FEP

In contrast to the optimal Bayesian update model, living animals cannot rapidly converge to the optimal posterior. We therefore assume that the agent updates its posterior gradually, following the free-energy principle (FEP). Within the FEP framework, the posterior distribution of *θ* is approximated by a Gamma distribution,

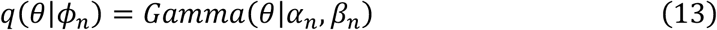

where *ψ_n_* = {*α_n_*, *β_n_*} denotes the shape and rate parameters at trial n, respectively. At each trial, the agent updates *ψ_n_* by minimizing surprise, defined as − log *p*(*o_n_*|*o*_1:*n*_). This quantity can be decomposed as

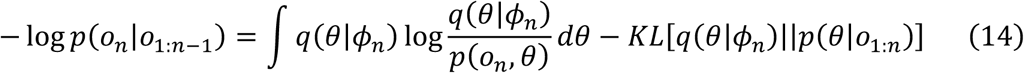

where *KL*[*q*(*x*)||*p*(*x*)] denotes the Kullback-Leibler divergence between distributions *q*(*x*) and *p*(*x*). Because the KL divergence is nonnegative, the first term provides an upper bound on surprise, known as the variational free energy,

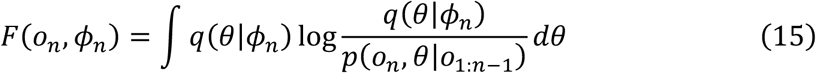

At each trial, the agent updates *φ_n_* by minimizing the free energy as,

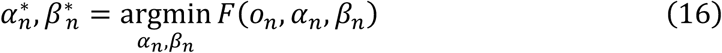

Because direct optimization over *α_n_* and *β_n_* may yield negative values, despite these parameters being strictly positive, we reparameterize them as *α_n_* = exp *u_n_* and *β_n_* = exp *v_n_* and perform optimization with respect to *u_n_* and *v_n_* . This leads to gradient descent updates of the form

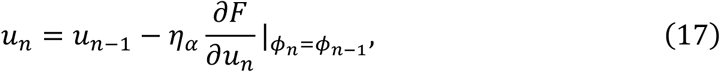

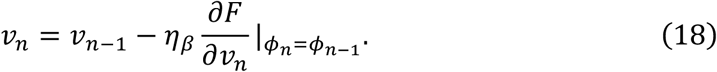

Transforming back to *α_n_* and *β_n_*, the updates become multiplicative,

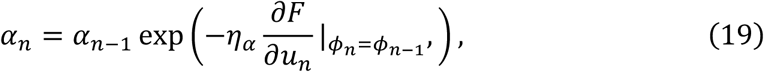

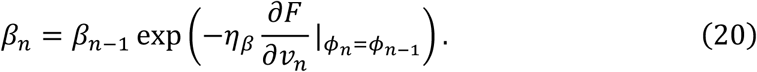

Using the chain rule, 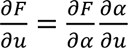 and 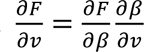, the gradients with respect to *α* and *β* are given by

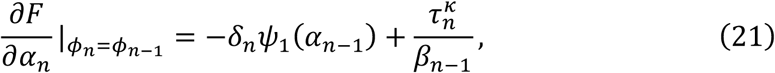

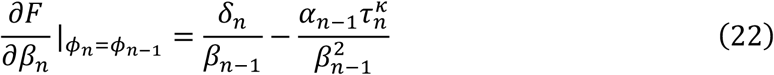

where *ψ*_1_(*x*) denotes the trigamma function. Substituting these expressions yields the final update equations,

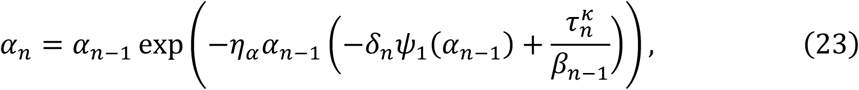

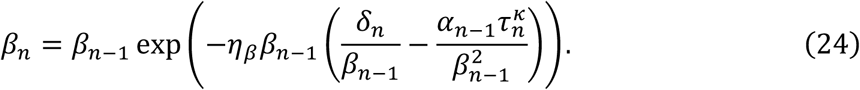

### Predictive distribution of explosion timing

Given the posterior distribution of *θ*, the predictive distribution of explosion timing on trial *n* is obtained by marginalizing *θ*:

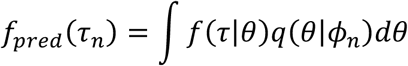

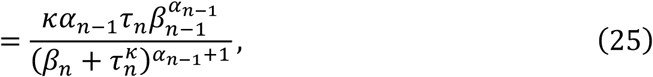

where *f*(*τ*) is the Weibull likelihood (Eq. (5)) and *q*(*θ* ∣ *φ _n_*_–1_) is the posterior belief from the previous trial.

### Expected reward

The expected reward is computed as a function of the cash-out time *T*. In the BART, the reward increases linearly with time if cash-out is successful, whereas no reward is obtained if an explosion occurs. The reward is therefore defined as

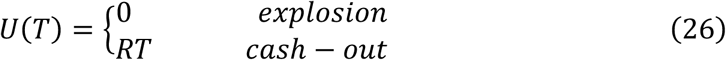

where *R* denotes the rate of reward increase over time. Thus, the expected reward at cash-out time *T* is given by

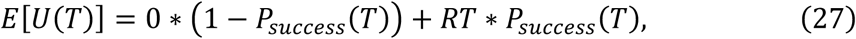

where *P_success_*(*T*) denotes probability that the agent successfully obtained the reward by the cash-out timing *T*. This success probability corresponds to the probability that no explosion has occurred up to time *T*, and is given by the predictive survival function,

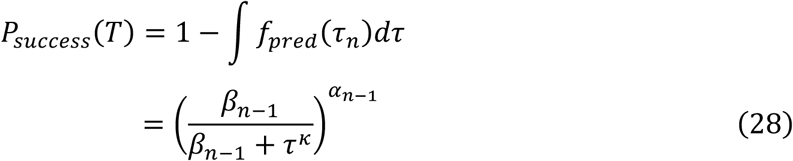

### Action selection depending on the optimism-pessimism bias

Based on the expected reward, the optimal cash-out timing is obtained by maximizing the expected reward with respect to the cash-out time *T_n_*:

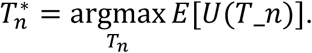

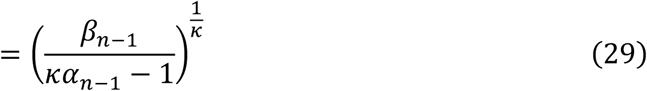

where *α_n_*_–1_ and *β_n_*_–1_ are the parameters of the Gamma distribution for the posterior at trial *n* − 1. We assumed that the agent has OP bias which distorts optimal cash-out timing as follows:

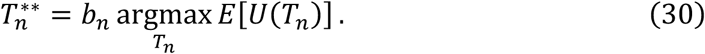

With this formulation, the agent can take risk-taking and risk-avoidance behaviors.

### Simulation protocol of the model agent

In simulations of the model agent, trial-by-trial changes in *α_n_*, *β_n_*, and actions were numerically calculated, *α_n_* and *β_n_* were updated by Eqs. (23) and (24), respectively. In each trial, the action was determined by Eq. (30) with the OP bias (*b_n_*) which was given arbitrarily (*b_n_* = 1 for **Fig. 3a**, b; *b_n_* = 1.3 for **Fig. 3d**; *b_n_* = 0.7 for **Fig. 3e**). In each trial, the either explosion or not was sampled from the Weibull distribution in Eq. (4).

### State-Space Model (SSM)

From the perspective of an external observer, the agent’s internal states are not directly accessible and must be estimated from observable behaviors. A state-space model (SSM) is therefore formulated to capture both the temporal evolution of latent internal states and the probabilistic generation of observable behavior from those states.

### Latent state dynamics

At trial *n*, the latent state is defined as *z_n_* = {*b_n_*, *α_n_*, *β_n_*}. Because no prior knowledge is assumed regarding the temporal evolution of the OP bias, it is modeled as a random walk. Since *b_n_* is constrained to be nonnegative, the random walk is modeled in log space:

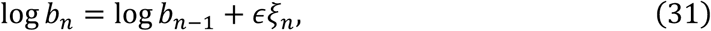

where *ξ_n_* ∼ *N*(*ξ_n_*|0, 1) is Gaussian noise and *ε* controls the magnitude of trial-to-trial fluctuations. The cognition parameters *α_n_* and *β_n_* evolved deterministically by the sub-optimal Bayesian update rules (Eqs. (23) and (24)).

### Observation model

The observable data consist of the waiting time until an event occurs on each trial (*τ_n_*), together with the event type (*δ_n_* = 1 for explosion, and 0 for successful cash-out). Thus, the observable data at trial n are defined as *o_n_* = (*δ_n_*, *τ_n_*). The cash-out timing *T_n_* is deterministically specified by the policy derived from the OP-modulated optimal timing (Eq. (30)). However, to improve numerical stability for inference, the observed cash-out time is assumed to follow a log-normal distribution:

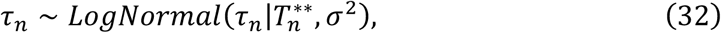

Where 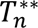 is the optimal timing (Eq. (30)), and *σ*^2^ denotes the variance. The likelihood probability of explosion at time *τ_n_* is described by

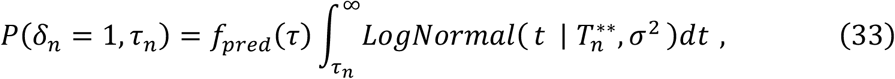

where *f_pred_*(*τ*) is the predictive distribution of explosion timing (Eq. (25)). On the other hand, the likelihood probability of cash-out at time *τ* is given by

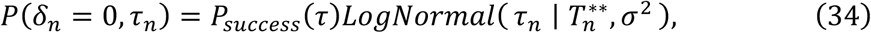

where *P_success_*(*T*) denotes the predictive survival probability up to time *T*.

### inverse OP by particle filter and smoother

Based on the SSM, the posterior distribution of the latent internal state of the agent *z_n_* = {*b_n_*, *α_n_*, *β_n_*} was estimated from behavioral observations, i.e., *P*(*z_n_*|*o*_1:*N*_) . This estimation was done by filtering and smoothing algorithms based on the Monte Carlo approximation^39^. In filtering, the posterior distribution of *z_n_* given observations up to trial *n* (*x*_1:*n*_) is sequentially updated in a forward direction as

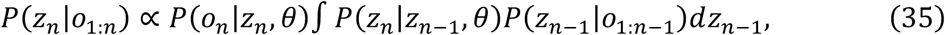

where *θ* = {*σ*^2^, *η_α_*, *η_β_*, *R*, *ε*}. The prior distribution of *b*_1_ is 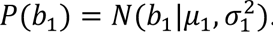. We used a particle filter to sequentially calculate the posterior *P*(*z_n_*|*o*_1:*n*_), which cannot be analytically derived because of the nonlinear transition probability. In our implementation, we adopted the Effective Sample Size (ESS) criterion to reduce computational cost of resampling^40^.

After the particle filter, the posterior *P*(*z_n_*|*o*_1:*N*_) is sequentially updated in a backward direction as

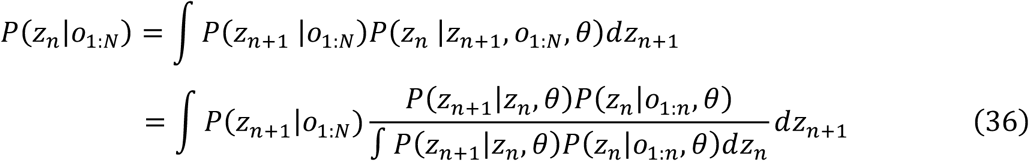

However, this backward integration is intractable because of the non-Gaussian *N*(*z*|*μ*, *σ*), which was represented by the particle ensemble in the particle filter, and the nonlinear relationship between *z_n_* and *z_n_*_+1_ in *P*(*z_n_*|*z_n_*_+1_). Then, a particle smoother was used (Forward Filtering-Backward Smoothing (FFBS)) to calculate the posterior *P*(*z_n_*|*o*_1:*N*_)^41^.

### Experimental protocol

One Japanese macaque monkey participated in this study (male, 5 years old, 9.0 kg). The monkey performed a modified version of the Balloon Analogue Risk Task (BART) for macaque monkeys, involving a trade-off between increasing reward and increasing risk of losing the accumulated reward.

At the beginning of each trial, two targets were presented on the screen: a central fixation target and a peripheral cash-out target located either to the left or right of the central target. The monkey was required to maintain fixation on the central target for 1.0 s to initiate the trial and to continue fixating on it while the balloon inflated. During this fixation period, a gray balloon-like stimulus centered on the central target inflated continuously, with its radius increasing at a rate of 0.7 deg/s. The size of the balloon corresponded to the amount of accumulated reward. The monkey could stop the balloon inflation by making a saccade to the peripheral cash-out target. If the saccade was made before the balloon exploded, the accumulated reward was secured and delivered.

Reward magnitude increased linearly as a function of inflation duration at a rate of 160 µL/s. Therefore, longer fixation resulted in a larger potential reward but also increased the probability that the balloon would explode. If the balloon exploded before the monkey made a saccade to the peripheral cash-out target, the trial was terminated and no reward was delivered.

Explosion times were predetermined to follow a uniform distribution between 2 and 6 s and were pseudo-randomly ordered across trials to reduce temporal bias.

## Author Contributions

I.H., K.-i.A., and H.N. conceived the project. I.H. and H.N. developed the computational framework. I.H. performed the analyses. R.I., R.Y., and K.-i.A. conducted the monkey experiments and acquired the data. Y.O. and K.I. contributed to refinement of the model and analysis framework. H.Y. and Y.Y. contributed to improvement of the manuscript and figures. H.N. supervised the study. All authors reviewed and approved the manuscript.

## Acknowledgements

This study was supported in part by the Moonshot R&D Program (JPMJMS2024-9 to H.N.), CREST (JPMJCR25Q2 to H.N.) from the Japan Science and Technology Agency (JST), Japan Agency for Medical Research and Development (AMED) Multidisciplinary Frontier Brain and Neuroscience Discoveries (Brain/MINDS 2.0) (JP25wm0625322 and JP25wm0625210 to H.N.), KAKENHI (JP21H03541 to H.N.), JST BOOST (JPMJBS2424 to I.H.), Naito Foundation (K.-i.A.), Takeda Science Foundation (K.-i.A.), Uehara Memorial Foundation (K.-i.A.), Japan Agency for Medical Research and Development JP24gm6910012 (K.-i.A.), JP24wm0625210 (K.-i.A.), JP21jm0210081 (K.-i.A.), Japan Society for the Promotion of Science JP24H02163 (K.-i.A.), JP21K19428 (K.-i.A.), JP21H05169 (K.-i.A.), JP20H03555 (K.-i.A.), JP20H05063 (K.-i.A.) JP22H04998 (K.-i.A.) and JP24KJ1424 (R.I.). Monkeys were provided by NBRP “Japanese Monkeys” through the National BioResource Project of the MEXT, Japan.

## Data availability

The data used in this study will be made publicly available on GitHub upon publication.

## Code availability

The code used in this study was implemented in Python (v3.11) and will be made publicly available on GitHub upon publication.

## Supplemental information

Supplemental information is available for this paper.

## Ethical approval

This study was conducted in accordance with the Guidelines for Care and Use of Nonhuman Primates, 3rd edition, and was approved by the Animal Experimental Committee of Kyoto University (approval number Med Kyo 25095). All animals were assessed to be in good health before participating in the study and had no prior experimental involvement.

---

**Supplementary Fig. 1:**
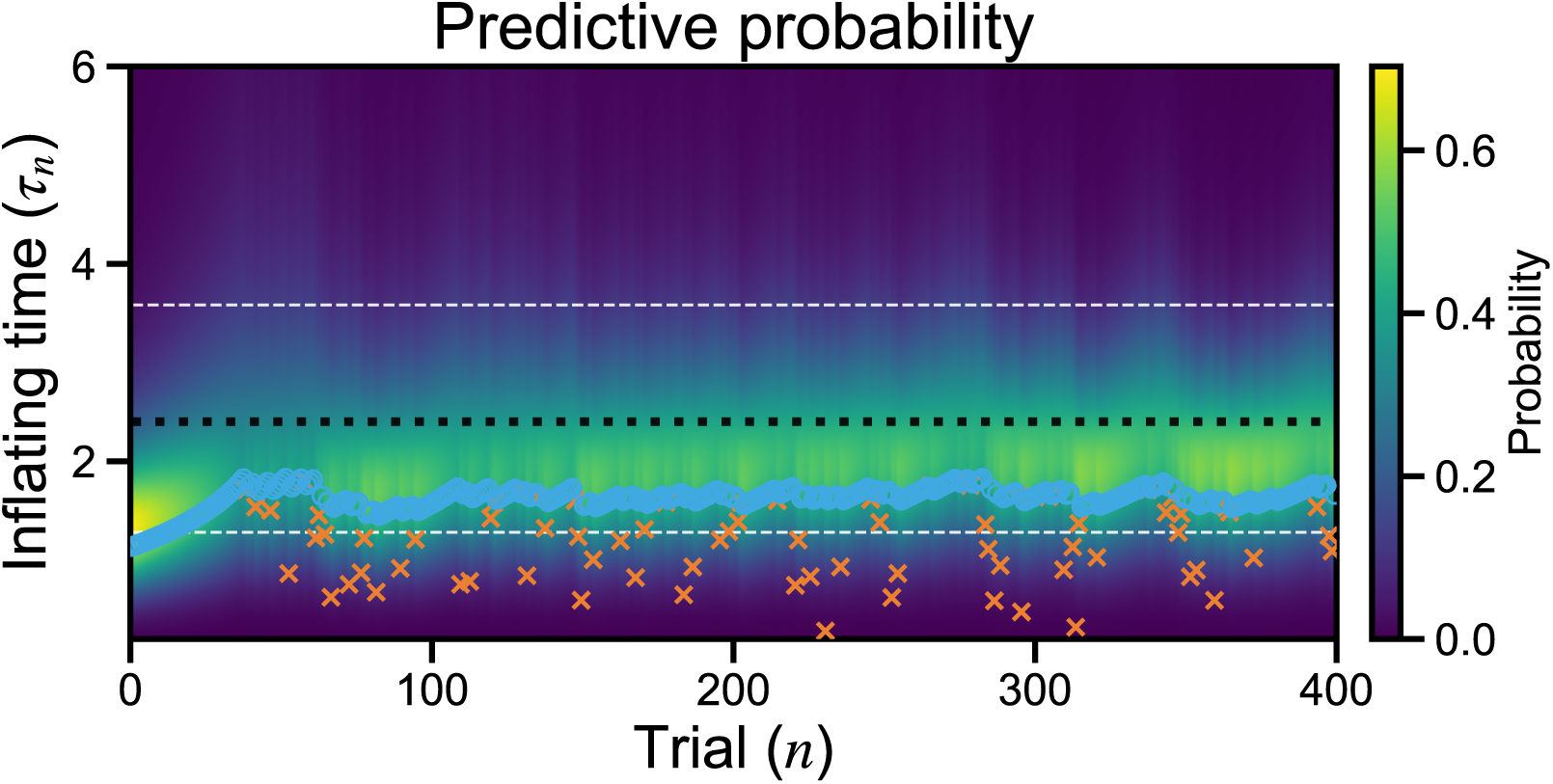
Predictive probability of overestimate. Predictive probability of explosion timing as a time-by-trial heat map with inconsistency of *κ* between agent and environment. *κ* = 4 for the agent and *κ* = 3 for the environment. Red cross and blue circle indicate timings of explosion and cash-out, respectively. Black and white dashed lines indicate median and 10%/90% quantiles of true explosion probability.

**Supplementary Fig. 2:**
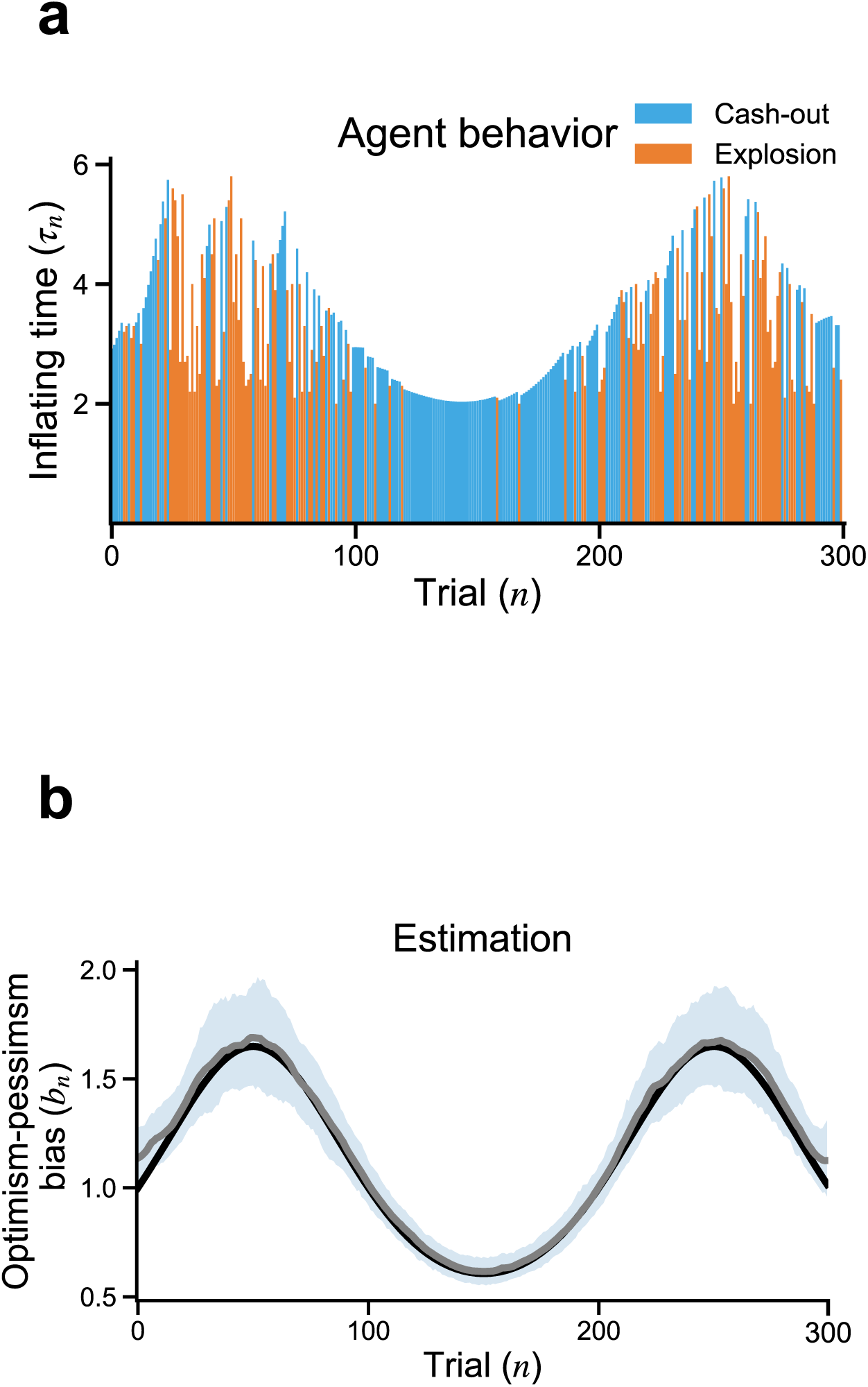
Trial-wise decoding of OP bias from artificial data. (**a**) Cash-out/explosion timings of the artificial agent. (**b**) inverse OP-decoded OP bias over trials. Black line is ground truth of OP bias. posterior mean (solid line) with 95% credible band (shading,2.5% 97.5%).

**Supplementary Fig. 3:**
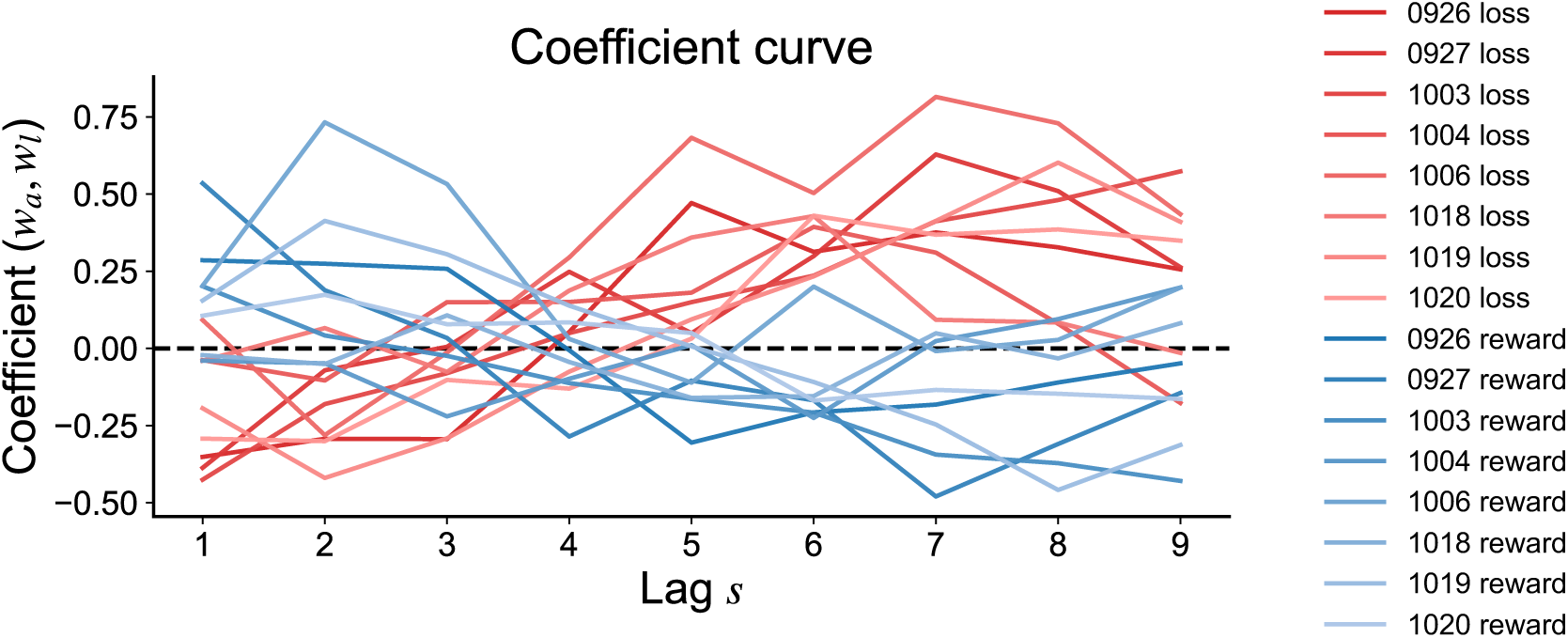
Coefficient curve of all days. Lag-wise coefficients of the logistic classifier. Blue and red lines indicate coefficients for reward and loss, respectively.

**Table 1.** Hyper parameter set Parameters used in particle filter and smoother. Some of these (*κ*, *ε*, *η_α_*, *η_β_*) were optimized using Bayesian optimization.

| Parameter | Value |
| --- | --- |
| $\kappa$ | 3.19182075203386 |
| $\epsilon$ | 0.09803506347411164 |
| $\eta_{\alpha}$ | 0.021348510178516772 |
| $\eta_{\beta}$ | 0.021447080785185497 |
| *particle # in paricle filter* | 1500 |
| $\sigma$ | 0.15 |
| *particle # in particle smoother* | 400 |
| *ESS* | 0.5 |
| $\alpha_1$ | 1.0 |
| $\beta_1$ | 50 |
| $\sigma_1$ | 0.2 |
| $\mu_1$ | 0.0 |
| $r$ | 1.0 |

